# Structured Gamma Spikes in Mouse Anterodorsal Thalamus

**DOI:** 10.1101/2025.02.22.639696

**Authors:** Tao Liu, Jia Liu

**Affiliations:** Tsinghua Laboratory of Brain and Intelligence, Tsinghua University, Beijing, China; Department of Psychological and Cognitive Sciences, Tsinghua University, Beijing, China

## Abstract

Gamma-frequency oscillations (∼30–160 Hz) are a hallmark of neuronal synchronization, yet the fine-scale temporal arrangement of spikes within individual gamma cycles remains poorly understood. Here, we examine head direction (HD) cells in the mouse anterodorsal thalamic nucleus (ADn)—a circuit distinguished by prominent high-gamma activity—to uncover the principles governing gamma coordination. We reveal two fundamental mechanisms: (i) a stable anatomical gradient in which neurons firing earlier in the gamma cycle exhibit longer anticipatory time intervals (ATIs), and (ii) a dynamic rate-phase shift whereby spike timing advances as the animal’s head aligns with a neuron’s preferred direction. Together, these results delineate a spatiotemporally structured framework for gamma synchronization, advancing our understanding of the functional roles and circuit mechanisms underlying gamma rhythms in local brain networks.

## 1 Introduction

Gamma-frequency oscillations are widely recognized as a fundamental mechanism for neuronal synchronization, appearing throughout mammalian brain regions—from the cerebral cortex (*1, 2*) and hippocampus (*3, 4*) to the olfactory bulb (*5, 6*) and thalamic nuclei (*7*). Emerging from local interactions between inhibitory interneurons and excitatory principal neurons (*8, 9*), these oscillations generate rhythmic patterns that regulate the timing of collective neuronal firing. Gamma rhythms have been proposed to bind distributed neuronal ensembles (*10*), enhance interregional communication (*11*), and support cognitive functions such as attention (*12, 13*), memory (*14, 15*), and sensory integration (*16*).

Although gamma oscillations primarily function to synchronize neural activity, systematic variations in spike timing are still evident across neurons and cognitive contexts (*17*). For theta oscillations (4–12 Hz), such temporal organization—exemplified by hippocampal place cell phase precession (*18*) and traveling waves (*19*)—has revealed key principles of information processing (*20*). By comparison, the detailed structure of gamma phase dynamics has been less extensively characterized [but see (*17, 21*)], and its functional role remains a subject of ongoing discussion (*22–24*).

The mouse ADn offers an ideal model to bridge our understanding of gamma phase dynamics. In contrast to other regions—where responses are dominated by external stimuli and multiple gamma sub-bands co-occur (*25, 26*)—ADn HD cells (*27*) encode a single intrinsic variable while exhibiting pronounced high-gamma oscillations (∼130–160 Hz; see (*28*)). Given the evolutionary conservation of brain architecture, the mechanisms revealed in the ADn are likely to extend to a broad array of regions exhibiting gamma oscillations.

Leveraging this simple and well-defined circuit, we demonstrate that gamma oscillations orchestrate HD cell firing with a robust temporal structure that persists across brain states yet is modulated by cognitive context. Specifically, through pairwise cross-correlation analyses and spike-triggered local field potential (LFP) phase measurements, we reveal that HD cells fire in a consistent, sequential order arranged both anatomically and functionally. Moreover, this sequential organization is dynamically adjusted in relation to individual neurons’ activation levels.

## Results

### HD Cells Identification

We analyzed data from a publicly available dataset (*29*). In this dataset, electrical signals were recorded from the mouse ADn during both free exploration and sleep using linear electrode arrays with 200 μm inter-shank spacing oriented perpendicular to the brain’s midline (Figure 1a). This configuration allowed us to simultaneously capture activity across distinct medial-lateral positions. HD cells were identified by applying a stringent criterion: a neuron was classified as an HD cell if its firing rate increased at least 10-fold when the animal’s head was aligned with its preferred direction compared with its least preferred direction (*30*) (Figure 1b). Across 25 recording sessions in six mice, we identified 183 HD cells (44.7% of recorded neurons), with most located on two adjacent electrode shanks (figure S1). We did not observe an overall difference in the mean firing rate of HD cells recorded from different shanks (figure S2).

**Figure 1:**
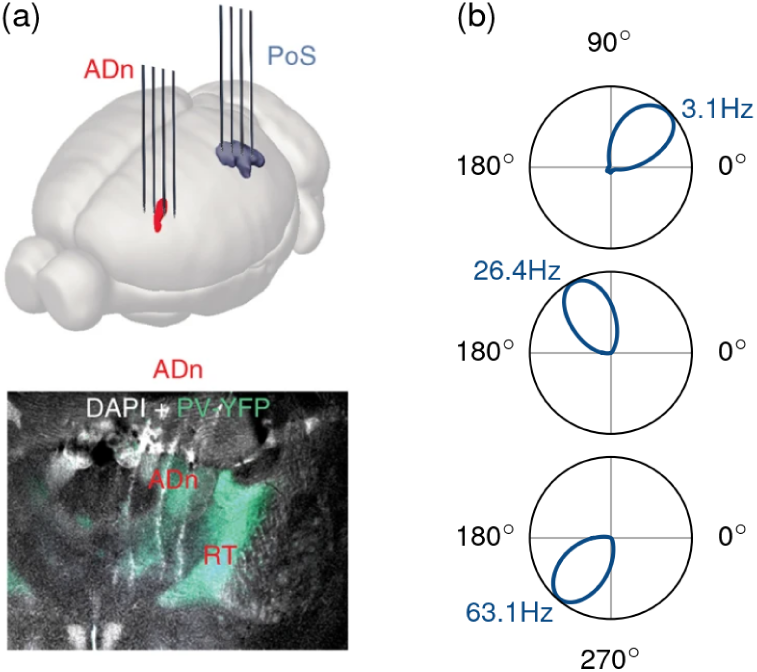
HD cells recorded in the mouse ADn. (**a**) Top: Schematic of the recording sites in the ADn (red). In some sessions, signals were also recorded from the post-subiculum (PoS, blue) but are not analyzed here. Bottom: A stained coronal section through the ADn. Panel (**a**) is adapted from (*28*). (**b**) Tuning curves of three representative HD cells. In each polar plot, the angular dimension represents head direction, and the radial dimension indicates the neuron’s mean firing rate. Peak firing rates (in Hz) are annotated at the maxima of the tuning curves.

### Gamma Synchrony Across Brain States

After establishing a robust population of HD cells, we probed how gamma oscillations orchestrate their activity by employing two complementary analyses: spike–spike cross-correlation and spike–LFP phase analysis. The former compares the firing times of neuronal pairs—revealing pair-specific synchrony—while the latter provides a global reference by aligning individual spikes to the phase of the population activity, albeit with signals that integrate inputs from diverse sources (*31,32*). Together, these approaches yield a comprehensive picture of gamma synchronization in the ADn.

We began by computing cross-correlation functions (CCFs) for neuronal pairs to assess their spike–spike relations. Traditional CCF methods, which bin spike trains, can introduce biases due to bin size selection. To overcome this, we developed a binning-free algorithm that computes CCFs with sub-millisecond precision [see (*30*) for details]. Focusing on pairs exhibiting a central peak within ±3 ms of 0 ms over a ±100 ms (or ±50 ms; see (*30*)) window, we found that 44.3% (101/228) of pairs displayed central peaks flanked by oscillatory sidebands—a clear signature of rhythmic, synchronized firing (Figure 2a). Importantly, these peaks were not strictly centered at zero; rather, each pair exhibited a characteristic phase offset that remained remarkably stable across wakefulness, REM sleep, and slow-wave sleep (SWS) (Figure 2a-c), underscoring the role of intrinsic local circuits in mediating this synchrony.

**Figure 2:**
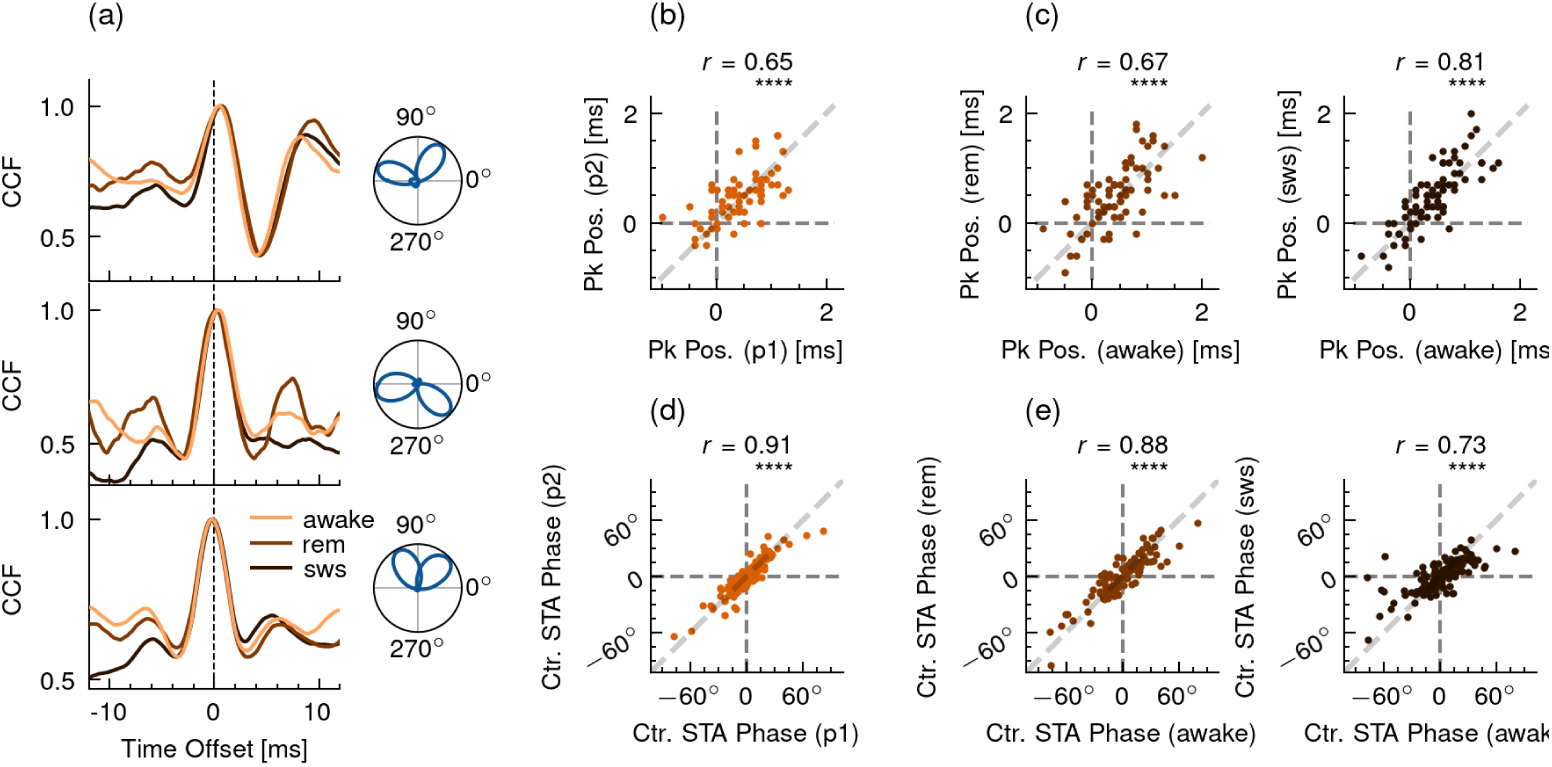
Synchronization of neuronal firing across brain states. (**a**) CCFs for three example neuronal pairs recorded during awake, REM, and SWS states (left). The awake-state CCF is normalized by dividing by its maximum amplitude. For visual comparison, the CCFs in REM and SWS are scaled to match the amplitude range of the awake-state CCF within ±5 ms. On the right are the normalized tuning curves of the same neuron pairs, illustrating each pair’s head-direction selectivity. (**b**) Peak positions (Pk Pos.) of the CCFs measured during two separate epochs of wakefulness, with each epoch’s data centered. (**c**) Peak positions of the CCFs during REM (left) and SWS (right) plotted against their corresponding peak positions during wakefulness. (**d**) STA phases of HD cells recorded during two separate epochs of wakefulness, with each epoch’s data centered (Ctr.). (**e**) STA phases of HD cells recorded during REM (left) and SWS (right) plotted against those in the awake state, with data centered within each state. In **b**-**e**, Pearson correlation tests were used to assess significance (****: *p* < 1 × 10^−4^), and the correlation coefficients (*r*) are indicated.

We next set up to examine the spike–LFP relationship. The LFP was band-pass filtered between 100 and 200 Hz and converted into its analytic signal to extract instantaneous phase values. Each spike was then assigned the corresponding phase of the analytic LFP at the time of occurrence. Despite the raw LFP lacking pronounced gamma spectral peaks in this band (figure S3), nearly all HD cells (97.3%, 178/183) exhibited significant phase coupling to the trough (180^◦^) of the LFP fluctuations (spike-LFP coupling; Rayleigh test, *p* < 1 × 10^−3^; Figure 3a,b). Furthermore, spike-triggered average (STA) phase analysis—by averaging LFP segments time-locked to spike events—confirmed that each neuron maintained a unique phase preference that was consistent across different brain states (Figure 2d-e).

**Figure 3:**
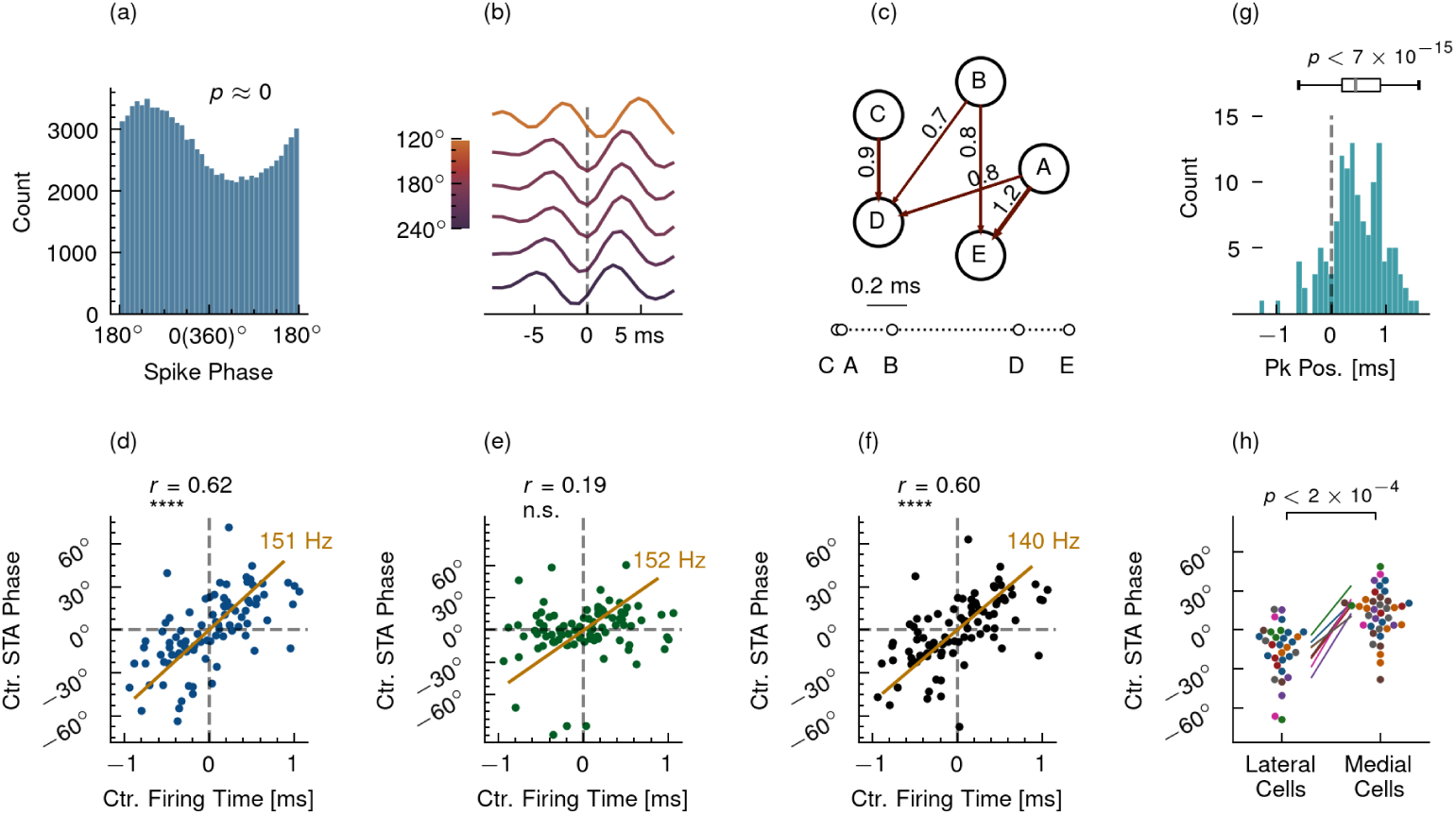
Firing sequences and anatomical gradient. (**a**) Distribution of spike phases for an example HD cell relative to 100–200 Hz LFP fluctuations, with a Rayleigh test assessing the significance of the phase distribution. Data in (**a**) are from session “Mouse25-140205”. (**b**) STA-LFP waveforms of HD cells from a single session (“Mouse17-130128”), normalized within ±8.8 ms, sorted by each cell’s STA phase, and displayed with artificial vertical spacing for clarity. (**c**) Top: Pairwise CCF peak positions among neurons, with arrows indicating which neuron leads. Only pairs exhibiting central peaks are shown. Bottom: The reconstructed firing sequence based on these offsets. Data in **c** are also from session “Mouse17-130128”. (**d**) Scatter plot of each neuron’s STA phase against its firing time in the reconstructed sequence, with values centered for each session. The orange line shows the orthogonal distance regression (ODR) fit, and the slope (151 Hz) is annotated. (**e**) Same as **c**, but the LFP was band-pass filtered in the 200–400 Hz band. (**f**) Same as **c**, but the LFP was recorded from shanks devoid of HD cells. In **d**-**f**, Pearson correlation tests were used to assess significance (n.s.: not significant, *p* > 0.05). (**g**) Histogram of CCF peak positions, taking medial spike trains as the reference. A binomial test was used to assess whether the distribution is significantly greater than zero. (**h**) Comparison of STA phases between lateral and medial HD cells, centered for each session. Mann–Whitney U rank tests were performed across eight sessions, and the resulting p-values were combined using Fisher’s method.

Collectively, these analyses demonstrate a robust and state-invariant temporal structure of HD cell firings in the ADn.

### Convergent Firing Sequences from CCFs and STA Phases

Both the pairwise phase offsets from our CCFs and the single-cell STA phases capture the fine-scale temporal structure of HD cell spiking. We asked whether these two measures would converge to reveal a consistent, population-wide sequential firing structure.

Specifically, the STA phase analysis estimates each neuron’s timing by determining the average gamma phase at which its spikes occur. Since spikes occurring earlier in the gamma cycle are associated with earlier phases, the STA phase naturally reflects a neuron’s position within the overall sequence (Figure 3b). In parallel, the CCF method quantifies the relative timing between neuron pairs: a positive peak indicates that the target neuron fires before its partner, whereas a negative peak denotes a delay. By aggregating these pairwise comparisons via regression [see (*30*) for details], we reconstructed a global firing order across the network (Figure 3c). As predicted, firing times derived from the CCF analysis correlated strongly with STA phase measurements (Figure 3d), underscoring the internal consistency of the sequential firing structure revealed by two methods.

Given the LFP reflects mixed aspects of population activity, we next asked which components of it do (or do not) contribute to the observed convergency of the two methodologies. We noted enhanced phase coupling of spikes to the LFP fluctuations in higher frequency bands—a phenomenon likely driven by residual spike waveform features remained in the LFP (*31*). In 56.3% of HD cells, the spike-LFP coupling was stronger in the 200–400 Hz band than in the 100–200 Hz band (103/183, based on Rayleigh test p-values). However, the correlation between CCF-derived firing times and STA phases was substantially lower for the 200–400 Hz oscillations (*r* = 0.19; Figure 3e) compared to the 100–200 Hz range (*r* = 0.62; Figure 3d). This indicates that while spike waveform contamination can affect the estimation of coupling strength, it does not preclude the extraction of meaningful timing information when focusing on an appropriate frequency band. Additionally, the convergence is unlikely to stem from the simple correlation of spikes with their own waveform residues in the LFP, as similar findings were obtained from LFPs that were recorded by shanks devoid of HD cells (Figure 3f).

Together, these findings demonstrate that the 100–200 Hz high-gamma oscillations provide an internal reference for the relative firing orders of HD cells.

### Anatomical Organization of Firing Sequences

We next investigated whether the structured firing of HD cells is anatomically organized within the ADn. Taking advantage of the electrode arrays that spanned the lateral-to-medial extent of the nucleus, we used each neuron’s shank location as an anatomical marker. Intriguingly, both the CCF-derived peak timings and STA phase measurements revealed a pronounced spatial gradient, with neurons on lateral shanks tended to fire earlier than those on medial shanks (Figure 2b-c, Figure 3g-h). On average, lateral neurons led their medial counterparts by 0.45 ms (from CCF peaks) and 37^◦^ (from STA phases). These observations indicate that gamma activity in the ADn propagates as a traveling wave from lateral to medial regions rather than emerging synchronously across the entire nucleus. Future studies with finer spatial resolution will help to further delineate the micro-architecture of this propagation.

### Firing Order Reveals a Functional Hierarchy

We next investigated whether the observed gradient in firing times plays a functional role in the HD circuit. Previous studies have linked spiking order to functional hierarchies across brain regions—for example, neurons in V1 fire earlier than those in latero-medial (LM) areas (*33*) and, along the HD pathway, ADn neurons precede those in the PoS (*28*). We therefore hypothesized that, within the region of ADn, neurons firing earlier in the gamma cycle act as upstream drivers for HD signal transmission.

To test this idea, we computed the ATIs (*34*) for individual HD cells (Figure 4a). The ATI quantifies how far in advance a neuron’s firing predicts the animal’s future head orientation, with larger ATIs implying a leading role in signal propagation (*35*). Across two distinct epochs in each recording session, HD cells exhibited highly consistent, cell-specific ATIs (Figure 4b), revealing an intrinsic temporal hierarchy in HD signal transmission.

**Figure 4:**
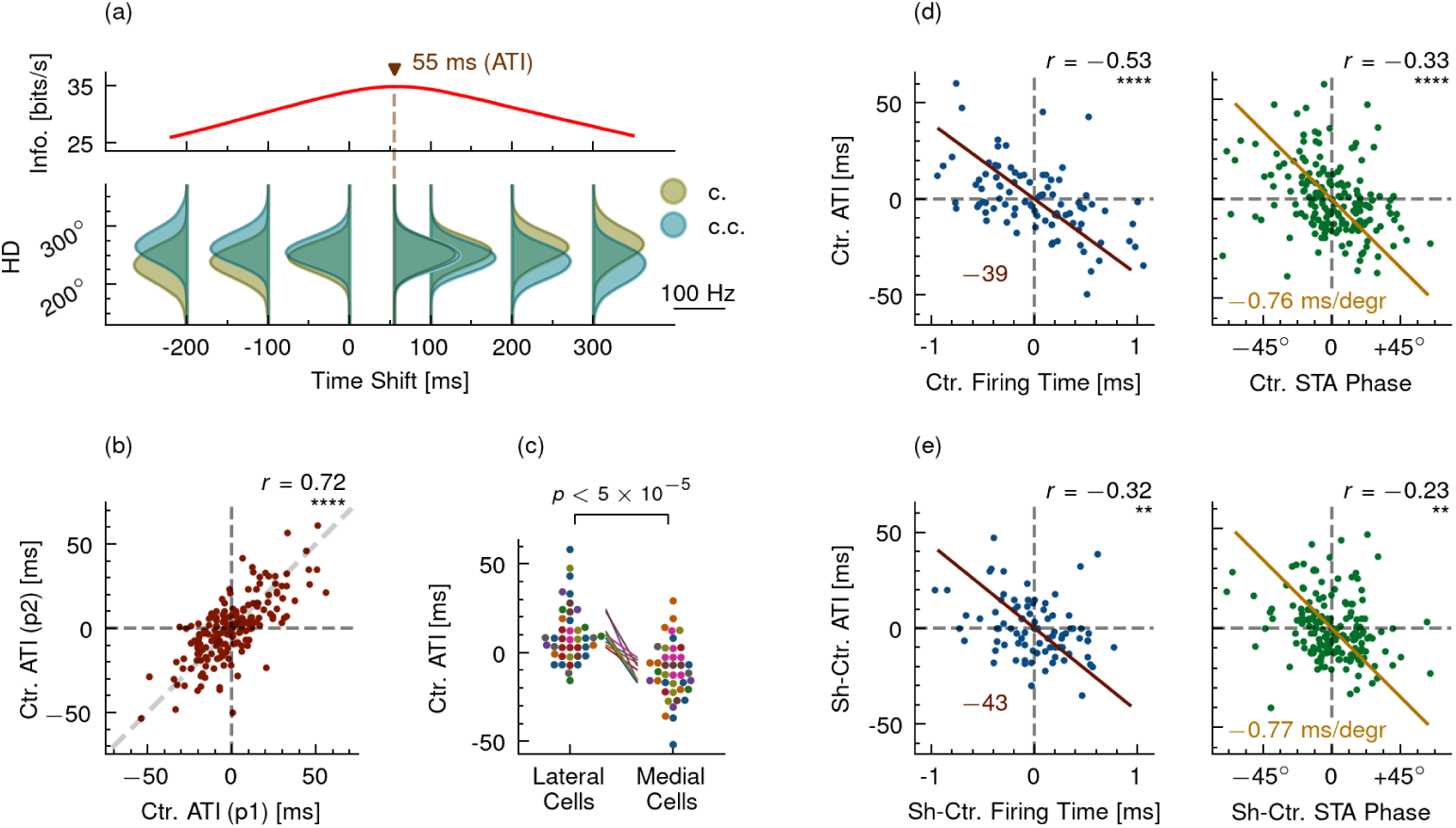
HD information flow aligned with spiking orders. (**a**) Top: Information content as a function of time shift for an example HD cell. The peak information content marks the ATI (≈55 ms here), which closely corresponds to the time shift where the cell’s clockwise (c.) and counterclockwise (c.c.) tuning curves diverge the least (*34*). Data in (**a**) are from session “B0703-211208” (*43*). Bottom: Clockwise and counterclockwise tuning curves at selected time-shift values for the same cell. (**b**) ATIs of HD cells measured during two separate epochs of wakefulness, with each epoch’s data centered. (**c**) ATIs of lateral versus medial HD cells, also centered by session. Mann–Whitney U rank tests were performed across nine sessions, and p-values were combined using Fisher’s method. (**d**) ATIs plotted against both reconstructed firing times (left) and STA phases (right), with data centered by session. The slopes of the ODR lines are annotated. (**e**) Same as (**d**), but with data centered within each shank group (Sh-Ctr.). Significance levels (**: *p* < 1 × 10^−2^) were assessed using Pearson correlation tests.

As hypothesized, we found that this functional hierarchy mirrors the anatomical gradient of firing order: HD cells recorded from lateral shanks displayed significantly higher ATIs than those from medial shanks, with a median difference of 30 ms (Figure 4c). Moreover, at the single-cell level, neurons that fired earlier within the gamma cycle tended to have higher ATIs compared to their later-firing counterparts (Figure 4d). This relationship persisted across different methods of sequence derivation and remained statistically significant after controlling for systematic shank-specific differences (Figure 4e).

Together, these results demonstrate that the sequential firing order of HD cells embodies a functional hierarchy that underpins the flow of directional information in the ADn.

### Dynamic Shifts in Firing Orders

While our earlier analyses demonstrated a robust and consistent firing sequence among HD cells, a closer look shows that this sequence is dynamically modulated by the animal’s cognitive contexts. In particular, the timing of spikes within each gamma cycle is systematically influenced by two interrelated factors: (i) the *directional deviation*—defined as the angular difference between the animal’s anticipated heading and a neuron’s preferred direction—and (ii) the neuron’s *firing rate quantile*, which reflects its instantaneous firing relative to its overall activity [see (*30*) for details]. Specifically, as the animal’s head more closely aligns with a neuron’s preferred direction, the neuron’s STA phase shifts to an earlier point in the gamma cycle (Figure 5a). Meanwhile, epochs of increased instantaneous firing are accompanied by an analogous phase advance—a pattern that holds across wakefulness, REM sleep, and SWS.

**Figure 5:**
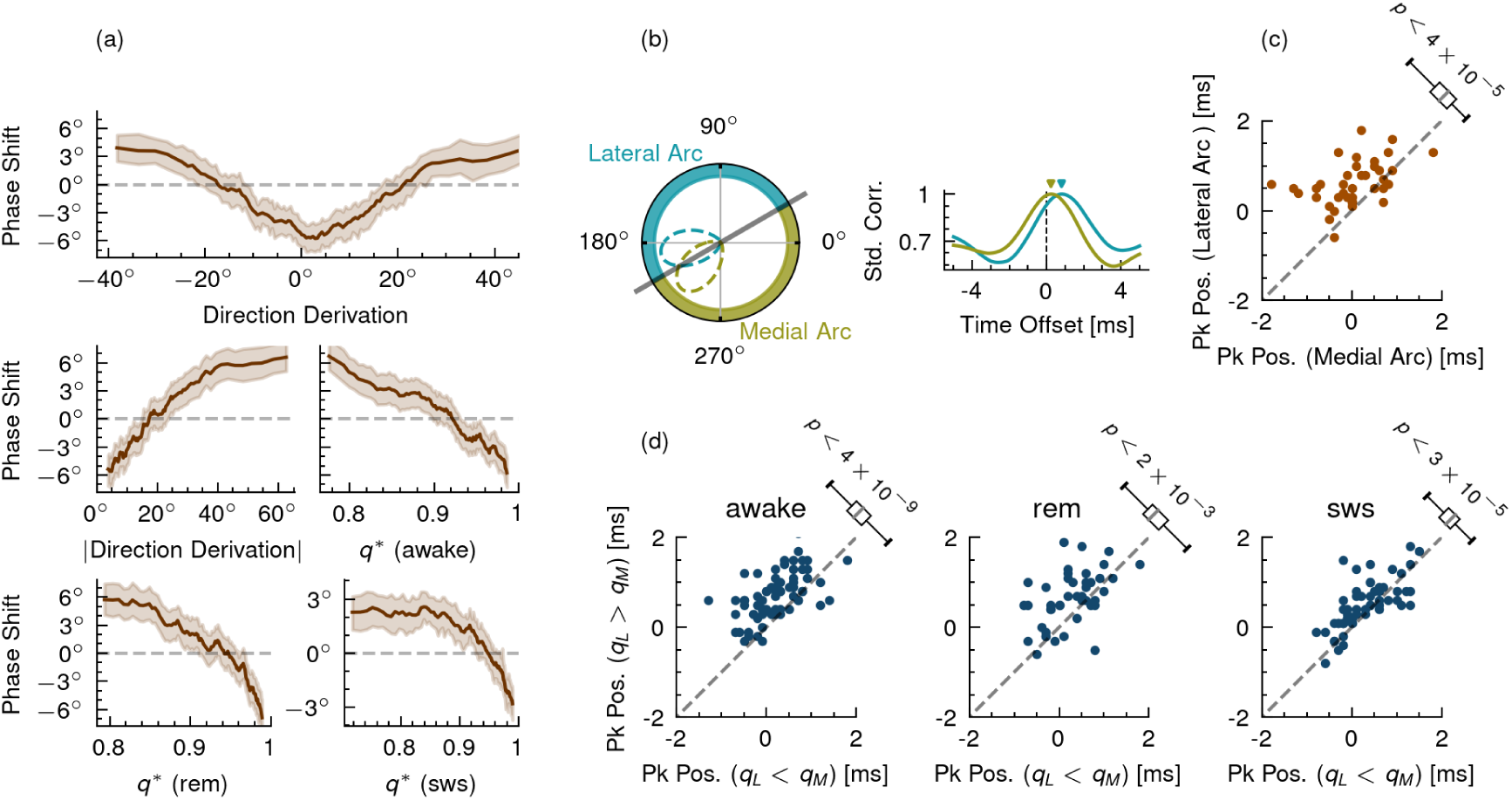
Advanced firing times when neurons prefer the heading. (**a**) Shift in STA phases as a function of head-direction derivation (top), the absolute value of direction derivation (middle-left), and firing rate quantile (*q*^∗^) (middle-right and bottom), shown across different brain states. Shaded areas indicate 95% confidence intervals. (**b**) Left: Segmentation of head-direction space into *Lateral* and *Medial Arcs* for an example neuronal pair, with the pair’s normalized tuning curves shown in dashed lines. Right: CCFs for the same pair when the animal’s heading falls within each arc. Data in (**b**) are from session “Mouse12-120807.” (**c**) Comparison of CCF peak positions for headings in the *Lateral Arc* versus the *Medial Arc*. A binomial test was used to assess statistical significance. (**d**) CCF peak positions when the lateral neuron’s firing rate quantile (*q*_*L*_) is higher than that of the medial neuron (*q*_*M*_) versus lower (*q*_*L*_ < *q*_*M*_), shown across different brain states. Binomial tests were performed to assess statistical significance.

Our spike–spike analysis further corroborates these observations. By partitioning the head-direction space into two 180° sectors—the *Lateral Arc* (head directions closer to the lateral cell’s preferred direction) and the *Medial Arc* (those closer to the medial cell’s; Figure 5b)—we observed significant shifts in CCF peak positions, with a median advance of 0.5 ms in the Lateral Arc relative to the Medial Arc (Figure 5c). In parallel, when we segmented the recording period based on whether the lateral cell’s firing rate quantile exceeded that of its medial partner, the corresponding CCF peaks shifted significantly (Figure 5d). Notably, the rate-based adjustments in phase offset were evident across brain states, with median shifts of 0.5 ms during wakefulness, 0.4 ms during REM sleep, and 0.2 ms during SWS.

Together, these findings demonstrate that the temporal order of HD cell firing is not fixed but is continuously modulated by ongoing behavioral and cognitive demands. In individual neurons, moments of increased instantaneous activity are linked to an advance in spike timing within the gamma cycle—a phenomenon that mirrors similar observations in the cerebral cortex [reported in (*17, 21*), but not in (*36*)].

## Discussion

Our study reveals a dynamic and structured organization of gamma-phase spiking in the mouse ADn. We show that HD cells preferentially fire at specific gamma phases, creating a sequential pattern that is both anatomically and functionally organized. This temporal structure is evident along a medial–lateral axis—with lateral cells firing earlier than medial ones—and is tightly correlated with ATIs and modulated by instantaneous firing levels. Moreover, this organized sequence persists across brain states including wakefulness, REM sleep, and SWS.

Our results do not support the notion of a fixed or precise spiking structure within gamma cycles that functions as a rigid coding scheme (*37*). Instead, they point to a stochastic structure that makes sense at the statistical or population level. Functionally, such an organization may serve to coordinate information processing within the HD circuit. In particular, HD cells that fire earlier in the gamma cycle—often exhibiting higher ATIs and closer alignment to the anticipated head orientation—may be optimally positioned to drive subsequent network activity (*38, 39*).

Moreover, the observed lateral–medial gradient in phase preferences implies an underlying asymmetry in ADn connectivity, raising intriguing questions about the roles of circuit architecture and spike timing–dependent plasticity (*40*) in establishing and maintaining this organization. Our findings also prompt further investigation into the topographical connectivity beyond the ADn along the HD pathway [see (*41*) for a review]. For instance, in the PoS—a downstream region that shares a common gamma clock (*28*) and is reciprocally connected with the ADn—anterior PoS primarily projects to the rostromedial ADn, whereas caudal PoS targets more rostrodorsal regions (*42*). Future studies are warranted to elucidate how such topographical connections contribute to the observed firing patterns.

Overall, by focusing on the ADn HD circuit that encapsulates key features of gamma synchronization, our findings offer a reductionist yet powerful framework for dissecting the circuit-level mechanisms underlying gamma-mediated neural timing and information processing. Future studies with enhanced spatial resolution, circuit mapping, and manipulation approaches will be essential to further elucidate how these temporal dynamics integrate with network connectivity and contribute to cognitive functions.

## Acknowledgments

Funding:

## Supplementary materials

Materials and Methods

Figs. S1 to S4

References (*42–46*)

## Materials and Methods

### Experiment

The experimental setup has been extensively described in (*28*), so here we only provide specifics pertinent to the context of this study. (i) To prevent statistical bias of neuron sampling, we exclusively utilized data from the first session whenever multiple sessions were conducted with electrodes maintained at identical locations. (ii) Neurons with an average firing rate below 0.5 Hz during awake states were excluded for analysis. (iii) Due to a random misalignment by 0-60 milliseconds in the first frame of the video used for extracting directional angles, we could not determine the non-relative values of the neuronal ATIs. Nevertheless, this misalignment did not affect the analysis of our study. For illustrative purposes, we used data from (*43*) to plot Figure 4a.

### HD Cell Identification

The metric employed to identify HD cells was the trough-to-peak ratio of their HD tuning curves. We found this ratio to be effective in distinguishing between neurons with pronounced versus weak HD tuning (figure S4). Only neurons exhibiting a trough-to-peak ratio less than 0.1 were identified as HD cells and selected for inclusion in subsequent analyses.

### Periods on Which Analyses Were Conducted

In this work, analysis was always conducted on the corresponding period. The period is defined primarily based on brain state—wakefulness (and two sub-epochs, obtained by evenly split the wakefulness period at a specific time point), REM sleep, and SWS—and additionally grouped into an overall “session period” (or “session”) that encompassed all three states and their transitions. Moreover, the analyses in Figure 4b and Figure 5c-d applied finer segments. By default, analyses were performed on the session period unless noted otherwise.

### Calculation of CCFs and Peak Positions

Analysis was constricted to neuron pairs recorded on different shanks. The medial shank spike train was designated as the reference and the lateral shank spike train as the target. Spike data were segmented by periods, and the CCF was calculated for each period.

#### Binning-free Algorithm

For each period, cross-spike intervals in the ±102 ms range were extracted and analyzed using kernel density estimation (KDE) with a 2 ms Epanechnikov kernel in a 0.1 ms step over a ±100 ms window. The resulting cross-spike-interval distribution was treated as the CCF.

To identify reliable central peaks, we observed that the maximum of the CCF within ±100 ms typically occurred within a narrow ±3 ms window (figure S5). In the wakefulness state, a central peak was declared if the CCF maximum within ±100 ms fell within ±3 ms. For REM, SWS, and the session period, a central peak was recognized only if (i) a central peak had already been identified in the corresponding wakefulness CCF for that neuron pair, and (ii) the maximum within a ±50 ms window of the current state’s CCF also fell within ±3 ms. This two-step procedure helps to exclude CCFs that may be obscured by noise. The narrower window for non-awake periods accounts for the rapid sweeping of internal heading during SWS (*28*) and meanwhile ensures a sufficient sample size for statistical testing. Only CCFs with central peaks in the corresponding periods were included in corresponding analyses.

### Reconstruction of Firing Sequence

In (*44*), a method was proposed to reconstruct the collective firing sequence based on the peak positions of pairwise CCFs, which requires that all pairs exhibit a central peak in their CCFs. In our experiment, however, most HD cells were recorded only from two shanks, and pairs lacking a reliable central peak were excluded from analysis. To address this limitation, we designed an algorithm that uses only those pairs of neurons with identified central CCF peaks for reconstruction. Specifically, neurons connected—either directly or indirectly—by these peak-based relationships were grouped into a connected component in a graph, and a least-squares approach was then applied to estimate the firing sequence based on declared peaks offsets. Although a single session could theoretically yield multiple disconnected graphs, this did not occur in our data.

### Computation of STA Phase

The LFP was obtained by averaging the EEG data recorded from all shanks implanted in the ADn. The LFP was downsampled to 1250 Hz, band-pass filtered using a rectangular filter in the 100–200 Hz range, and then transformed into its analytic representation via the Fourier transform by doubling the positive frequency spectrum and discarding the negative frequencies. This analytic signal was used to compute the phase for each spike and the STA phase for each neuron.

### Outliers Related to STA Phase Measurement

Neurons with STA phases that deviated substantially from the polar mean phase of simultaneously recorded cells (i.e., more than three standard deviations from the population distribution) were classified as outliers. Specifically, 2 out of 183 neurons in Figure 2d, 3/183 in Figure 2e (left), 3/183 in Figure 2e (right), 2/183 in Figure 3e-f, 2/114 in Figure 3h, 2/183 in Figure 4d (right), and 2/183 in Figure 4e (right) were identified as outliers. The inclusion or exclusion of these outliers does not affect any conclusions of this study.

### Computation of ATI

We estimated the ATI of HD cells using the method adopted in (*45*). In brief, angle data timestamps were shifted in 5 ms steps within a ±800 ms window, and a tuning curve was generated at each shift using 6° bins. We then calculated the information content associated with each time shift using the formula from (*46*). The resulting information-content-over-time-shift profile was smoothed using a 100 ms Gaussian kernel, and the time shift corresponding to the global maximum of the smoothed profile was taken as the neuron’s ATI.

### Sessions Selected for Statistical Testing

For analyses in Figure 3h, figure S2, and Figure 4c, we compared the STA phase, the mean firing rate, and the ATI for neurons recorded from different shanks, respectively. To ensure sufficient statistical significance of Mann-Whitney U rank test, we only included sessions in which each shank had at least two HD cells (not outliers).

### Computation of Firing Rate Quantile

The spike train was convert to the series of instantaneous firing rate through KDE estimation with a 100 ms Gaussian kernel and a 5 ms convolving step. Then, for each period, the instantaneous firing rate in the series was converted to the quantile value corresponding to its rank in this series.

---

**Figure S1:**
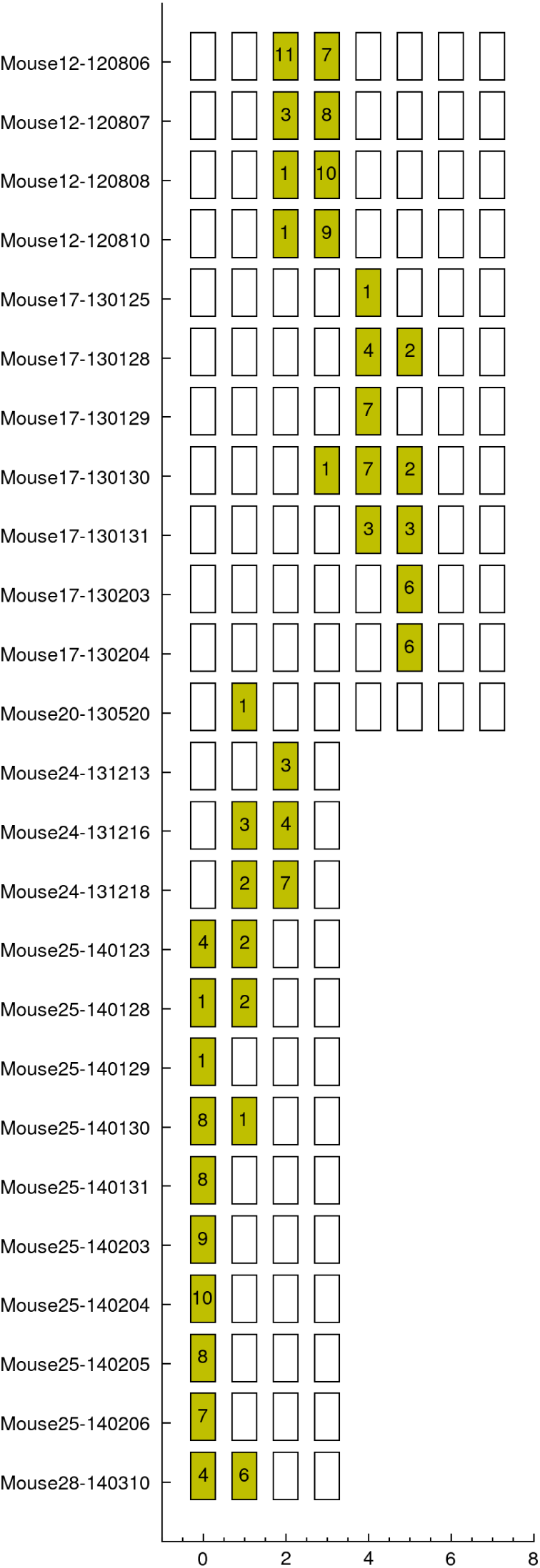
Distribution of HD cells across recording shanks. The figure displays the linear arrangement of recording shanks annotated with the number of HD cells recorded on each shank (ordered from lateral to medial). Notably, the majority of HD cells were clustered on two adjacent shanks.

**Figure S2:**
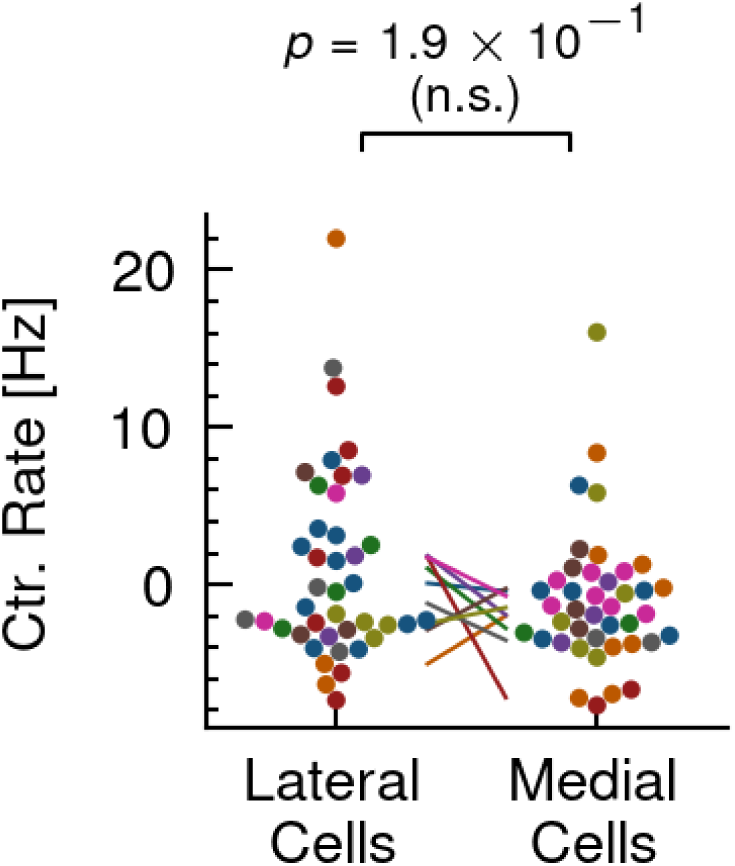
Firing rates of HD cells recorded from different shanks. Comparison of mean firing rates between lateral and medial HD cells, centered for each session. A two-sided Mann–Whitney U rank tests was performed. (n.s.: not significant, *p* > 0.05).

**Figure S3:**
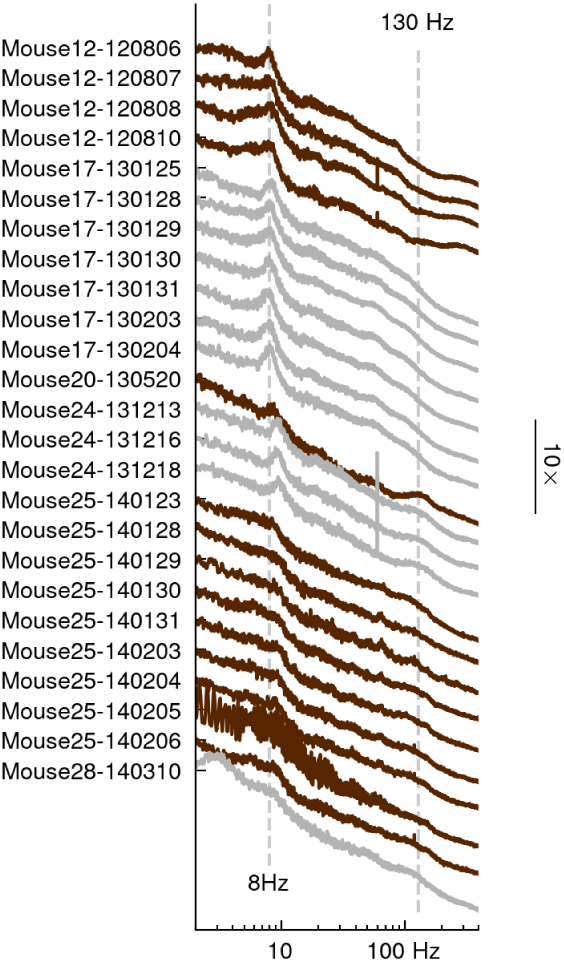
LFP Power Spectrum. Power spectra of the LFPs during wakefulness are shown, with curves vertically offset for clarity. While a prominent theta peak at 8 Hz is evident, no gamma peak (around 130 Hz) is observed in the raw LFP recordings.

**Figure S4:**
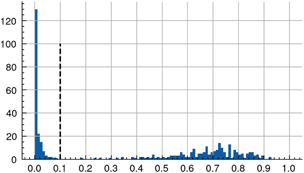
Distribution of trough-to-peak ratios in tuning curves. The histogram displays the distribution of trough-to-peak ratios computed from the tuning curves of all recorded neurons. Neurons with a trough-to-peak ratio below 0.1 were classified as HD cells. A clear boundary is evident in the distribution, distinguishing neurons with strong HD tuning from those with weak or absent tuning.

**Figure S5:**
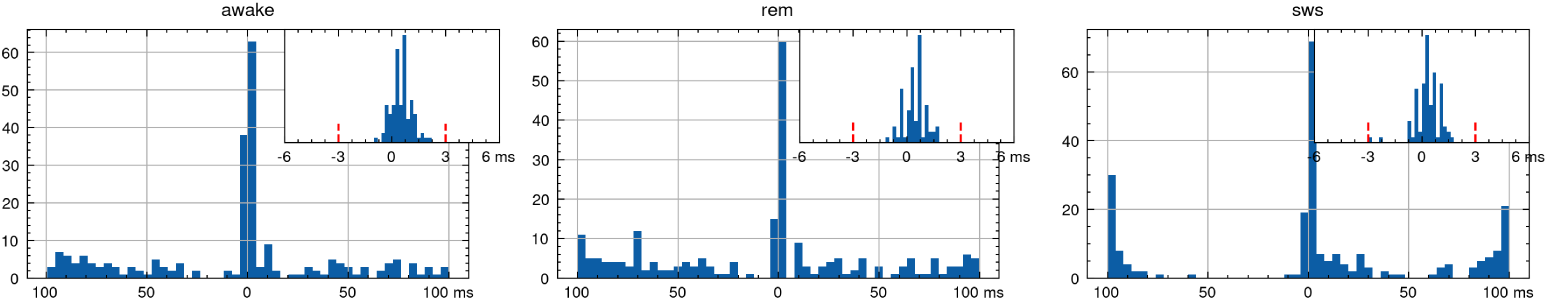
Distributions of CCF peak positions across brain states. Histograms display the distribution of CCF peak positions within a ±100 ms window during wakefulness, REM sleep, and SWS. An inset zoom focuses on the ±6 ms window. While peak positions during wakefulness, REM and SWS are concentrated within ±3 ms, the SWS distribution exhibits an increased occurrence of peaks extending beyond ±50 ms.

